## Supplementary Figures for "A conserved population of MHC II-restricted, innate-like, commensal-reactive T cells in the gut of humans and mice"

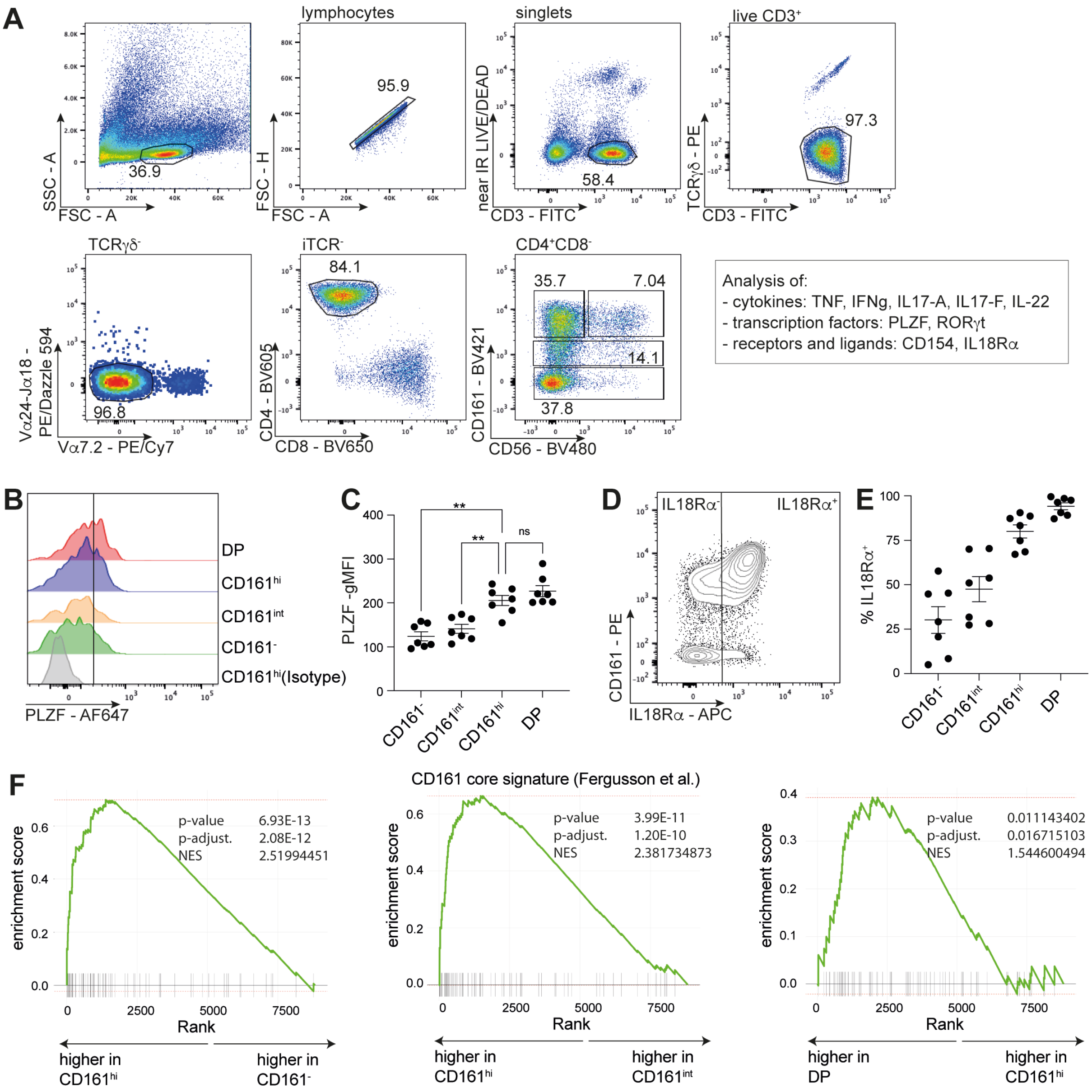


**Supplemental Figure 1: Human colonic CD161$^{hi}$ CD4 T cells show an enrichment of innate-like features and express a diverse TCR-repertoire. Related to Figure 1.**

(A) Gating strategy used to identify human colonic CD4 T cells not expressing TCRs associated with established innate-like T cell populations. Downstream analyses were performed on the four displayed CD4 T cell subsets or the entire CD4$^{+}$CD8$^{-}$ population. (B) Histograms derived from flow cytometric data summarizing the expression of the transcription and PLZF by the indicated CD4 populations. (C) Scatterplot showing the expression of PLZF as measured by gMFI on the indicated cell populations. (D) Flow cytometric analysis depicting the expression of IL18Rα in relation to CD161. The vertical line represents the cut-off used to define IL18Rα-positive cells. (E) Scatterplot illustrating the percentage of CD4 T cells from the indicated populations expressing IL18Rα. (F) CD161$^{-}$, CD161$^{int}$, CD161$^{hi}$ and DP CD4 T cells were sorted from freshly isolated LPMCs from three different individuals in bulk and subjected to RNA-sequencing. Gene set enrichment analyses were performed to assess the enrichment of a set of genes expressed in innate-like T cells and correlating with innate-like T cell responses (CD161 core, Fergusson et al, 2014) in the different CD4 subsets. The depicted GSEA plots show the comparisons of the CD161$^{hi}$ with the other three subsets. (C, E), Data points were pooled from independent experiments using one or two human samples each.. ns = not significant, **p < 0.01, repeated measures ANOVA with Tuckey's multiple comparisons test (C). Mean ± SEM is shown.

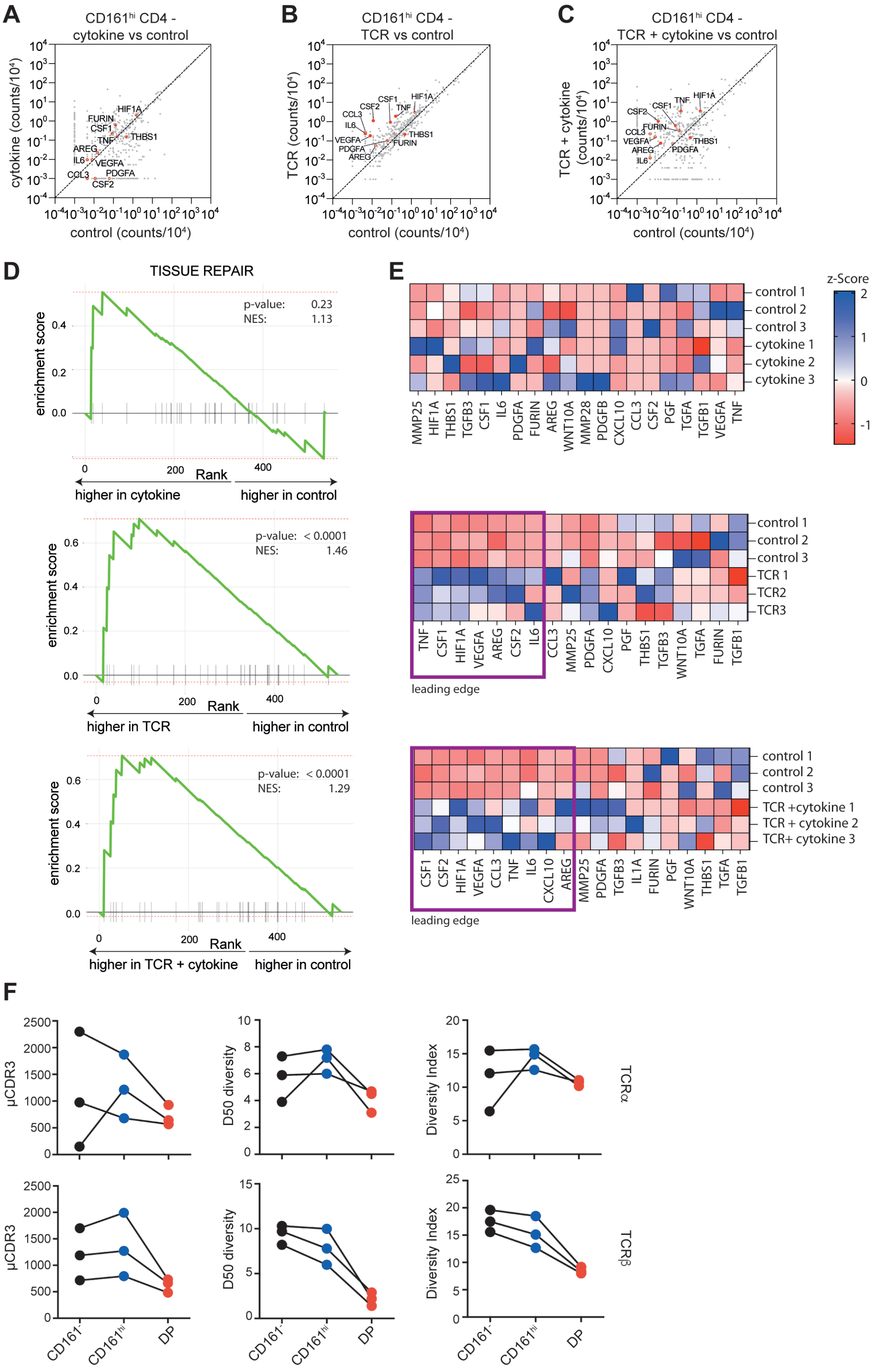

**Supplemental Figure 2: CD161$^{hi}$ CD4 T cells tissue repair-associated genes in a TCR-dependent manner and express a diverse TCR-repertoire. Related to Figure 1.**

(A-E) LPMCs from three donors were stimulated overnight with cytokines (IL-12+IL-18(=), plate-bound αCD3 antibodies or a combination of both, labelled with barcoded hashing and surface antibodies as well as fluorescently labelled antibodies for sorting. CD4 T cells negative for TCRs associated with established innate-like T cell populations were sorted and subjected to the BD Rhapsody single cell pipeline. CD161$^{hi}$ CD4 T cells were identified using the signal from a barcoded anti-CD161 antibody. (A-C) Diagonal plots showing the gene counts for individual genes found in the different stimulated versus the unstimulated CD161$^{hi}$ CD4 T cells. Genes from the tissue repair gene signature published by Linehan et al (2018) are marked in red. (D) GSEA analysing the enrichment of the tissue repair gene signature in CD161$^{hi}$ CD4 T cells stimulated with cytokines, plate-bound $\alpha$CD3 antibodies or both compared to unstimulated control cells. (E) Z-scores of the expression of the different genes from the tissue repair signature in CD161$^{hi}$ CD4 T cells from the individual donors. Leading edge genes driving the enrichment in (D) in the TCR and TCR + cytokine stimulated cells are marked. (F) Connected scatter plots showing the number of unique CDR3 regions, the D50 diversity and the Diversity Index calculated of the TCR$\alpha$ and $\beta$ chains used by FACS-sorted CD161$^{-}$, CD161$^{hi}$ or DP CD4 T cells. Both experiments were done with three donors.

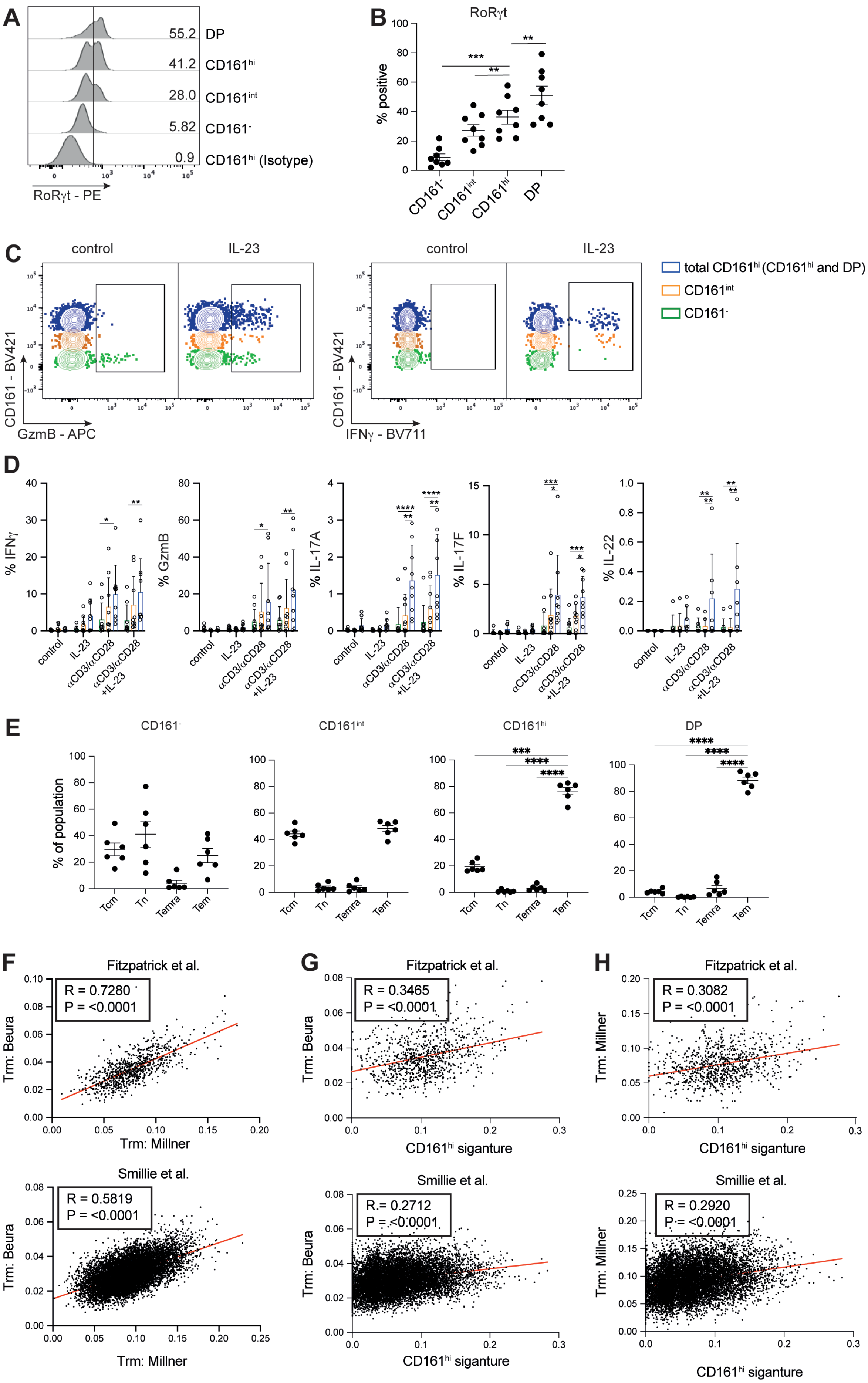

A
55.2 DP
41.2 CD161hi
28.0 CD161int
5.82 CD161-
0.9 CD161hi (Isotype)
RoRγt - PE
B
RoRγt
% positive
CD161-
CD161int
CD161hi
DP
C
control
IL-23
CD161 - BV421
GzmB - APC
IFNγ - BV711
total CD161hi (CD161hi and DP)
CD161int
CD161-
D
% IFNγ
% GzmB
% IL-17A
% IL-17F
% IL-22
control
IL-23
αCD3/αCD28
αCD3/αCD28 +IL-23
E
CD161-
CD161int
CD161hi
DP
% of population
Tcm
Tn
Temra
Tem
F
Fitzpatrick et al.
R = 0.7280
P = <0.0001
Trm: Beura
Trm: Millner
Smillie et al.
R = 0.5819
P = <0.0001
G
Fitzpatrick et al.
R = 0.3465
P = <0.0001
CD161hi siganture
Smillie et al.
R = 0.2712
P = <0.0001
H
Fitzpatrick et al.
R = 0.3082
P = <0.0001
Trm: Millner
CD161hi siganture
Smillie et al.
R = 0.2920
P = <0.0001

**Supplemental Figure 3: Colonic CD161$^{hi}$ CD4 T cells possess limited Th17 functionality, display a Tem phenotype while expression of their signature genes shows only a weak correlation with established $T_{RM}$ signatures. Related to Figure 1.**

(A, B) Histogramms derived from flow cytometric data and scatterplots summarizing the expression of the transcription factors RoR$\gamma$t by the indicated CD4 populations. (B) Flow cytometric data depicting the expression of GzmB or IFN$\gamma$ by colonic CD161$^{-}$, CD161$^{int}$, CD161$^{hi}$ or DP CD4 T cells after 20 hours of in vitro stimulation with IL-23. (C) Barplots summarizing the percentages of CD4T cells from the indicated populations expressing IFN$\gamma$, GzmB, IL-17F, IL-17A or IL-22 respectively after 20 hours of *in vitro* stimulation with recombinant IL-23, plate-bound $\alpha$CD3-antibodies or a combination of both.(E) Scatterplots summarizing the percentages of CD4 T cells from the indicated subsets displaying a naïve or either of the different memory phenotypes.Tn = naïve T cells (CCR7$^{+}$CD45RA$^{+}$), Temra = effector memory T cells re-expressing CD45RA (CCR7$^{-}$CD45RA$^{+}$), Tem = effector memory T cells (CCR7$^{-}$CD45RA$^{-}$) and Tcm = central memory T cells (CCR7$^{+}$CD45RA$^{-}$).(F – H) CD4 T cell populations were extracted from single-cell RNAseq datasets previously published by Smillie *et al*. and Fitzpatrick *et al*. Using the AUCell R package, the expression of two different $T_{RM}$ signatures, published by Milner *et al*. and Beura *et al*., and the expression of a 26-gene signature module obtained by comparing human colonic CD161$^{hi}$ and DP CD4 T cells to CD161$^{-}$ CD4 T cells was assessed in each single cell. The plots show the Pearson correlation of the two different $T_{RM}$ signatures and the CD161$^{hi}$ signature with either the Beura- or Milner- $T_{RM}$ signature in the two different single cell datasets. Data points were pooled from independent experiments using one or two human samples each

***$p < 0.001$, ****$p < 0.0001$; repeated measures ANOVA with Tuckey's multiple comparisons test (B, E). 2$^{nd}$-way ANOVA with Tuckey's multiple comparisons test (D), Mean ± SEM is shown.

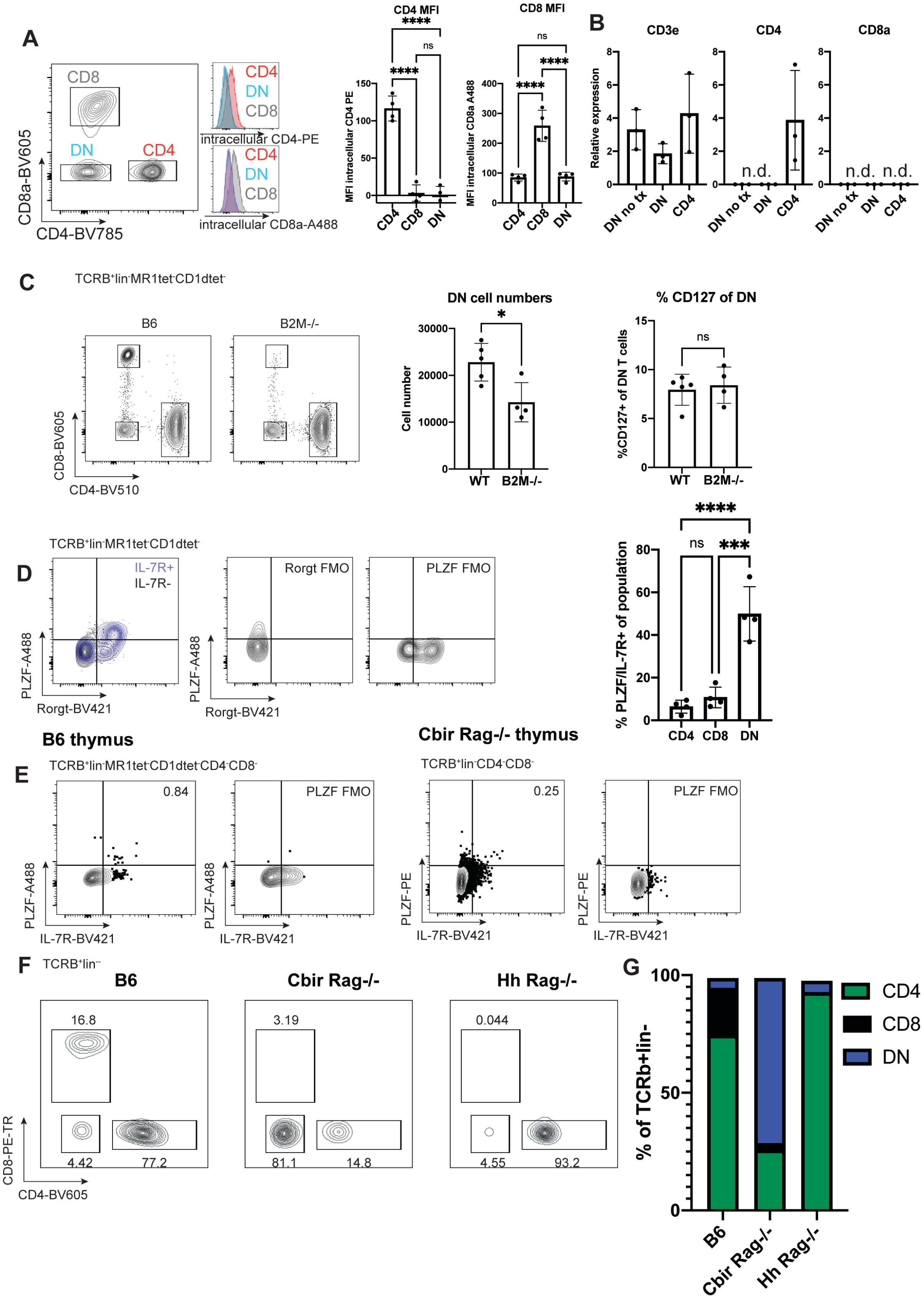


**Supplemental Figure 4: T$_{MIC}$ in mice have genuine lack of CD4 and CD8a, and requires microbial stimulus to develop**

(A) Representative flow cytometry (left) and quantification of mean fluorescence intensity (right) show intracellular CD4 stain in the CD4 population but not DN or CD8a populations from the MLN of Cbir Rag-/- mice. (B) qPCR of sorted colonic DN and CD4 T cell populations for *Cd3e*, *CD4* and *CD8* showing relative expression compared to *Hprt*. (C) Example flow cytometry (left) of gated TCRb+lin-MR1tet-CD1dtet- colonic LPLs from B6 and B2M-/- mice with quantification of DN cell numbers and the percent of the DN population with the IL-7R+ phenotype. (D) Representative FACS plots (left) show PLZF/RORgt expression (compared with FMOs) in the IL-7R+ population of DN T cells from B6 mice. Quantification of of PLZF/IL-7R+ proportion of CD4, CD8, and DN populations (right). (E) Representative flow cytometry plots of CD4, CD8a, and DN T cell populations in colonic LPL from B6, Cbir Rag-/-, and Hh7-2 Rag-/- mice (left) with quantification of the populations across the three genotypes.

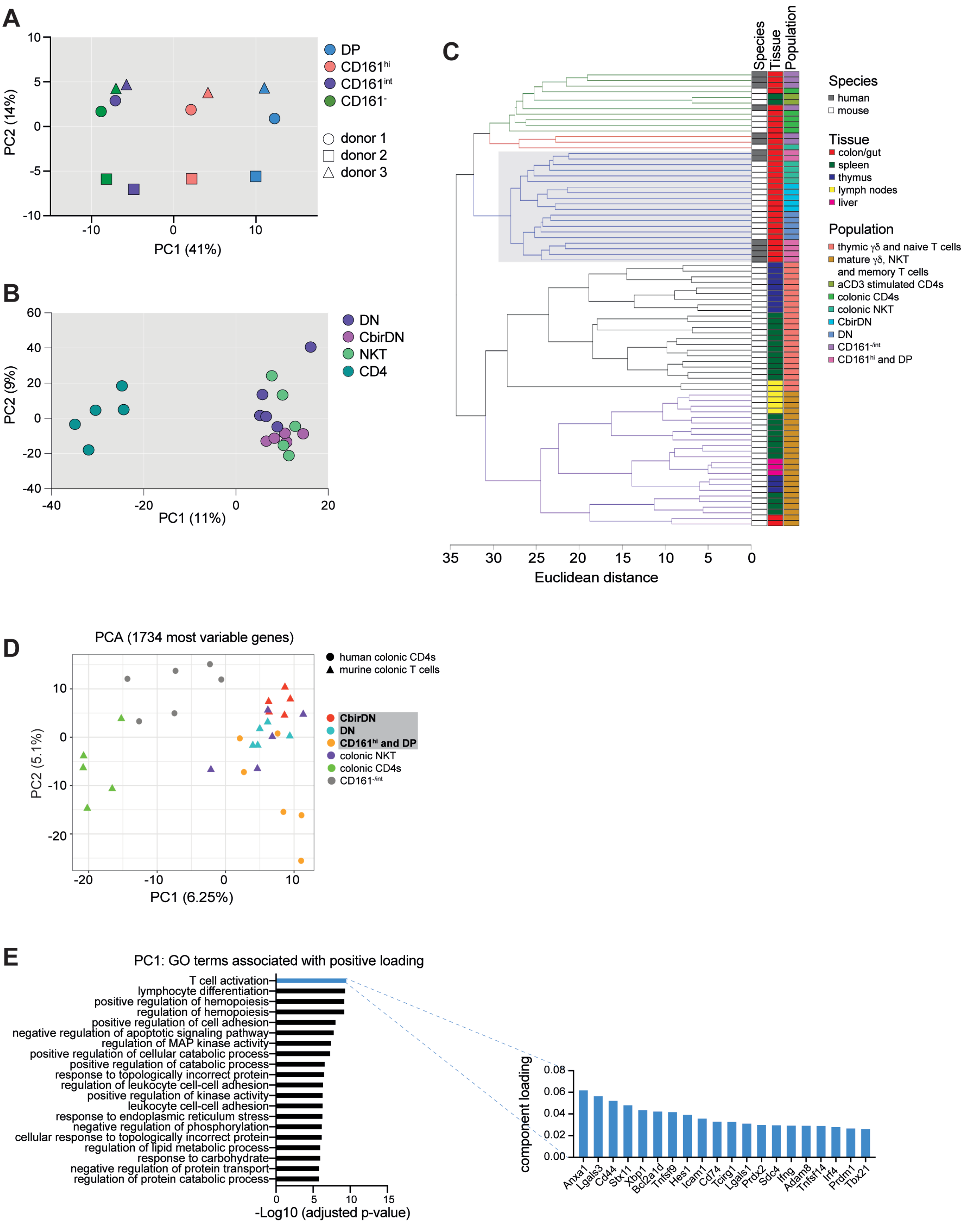


**Supplemental Figure 5: Transcriptional analysis of the separate and merged human and murine RNAseq-datasets. Related to Figure 4.**

(A, B) Principal component analyses performed on the genes expressed in bulk-sorted colonic human or murine T cell populations. The first two principal components are plotted.

(C) Clustering of the human and murine cells from (A, B) as well as several Immgen-derived RNAseq-datasets. To assess similarity, the Euclidean distance was computed based on the expression of the

866 most variable genes (IQR>0.75) derived from the merged dataset containing all the aforementioned datasets.

(D) PCA showing the clustering of the murine and human colonic T cell subsets from Figure 4 based on the most variable genes.

(E) Gene Ontology (GO) term enrichment analysis showing the top 20 GO-term enriched in the genes positively contributing to PC1 from the merged dataset from (C). For the highlighted GO-term, the corresponding top 20 genes from PC1 are shown.

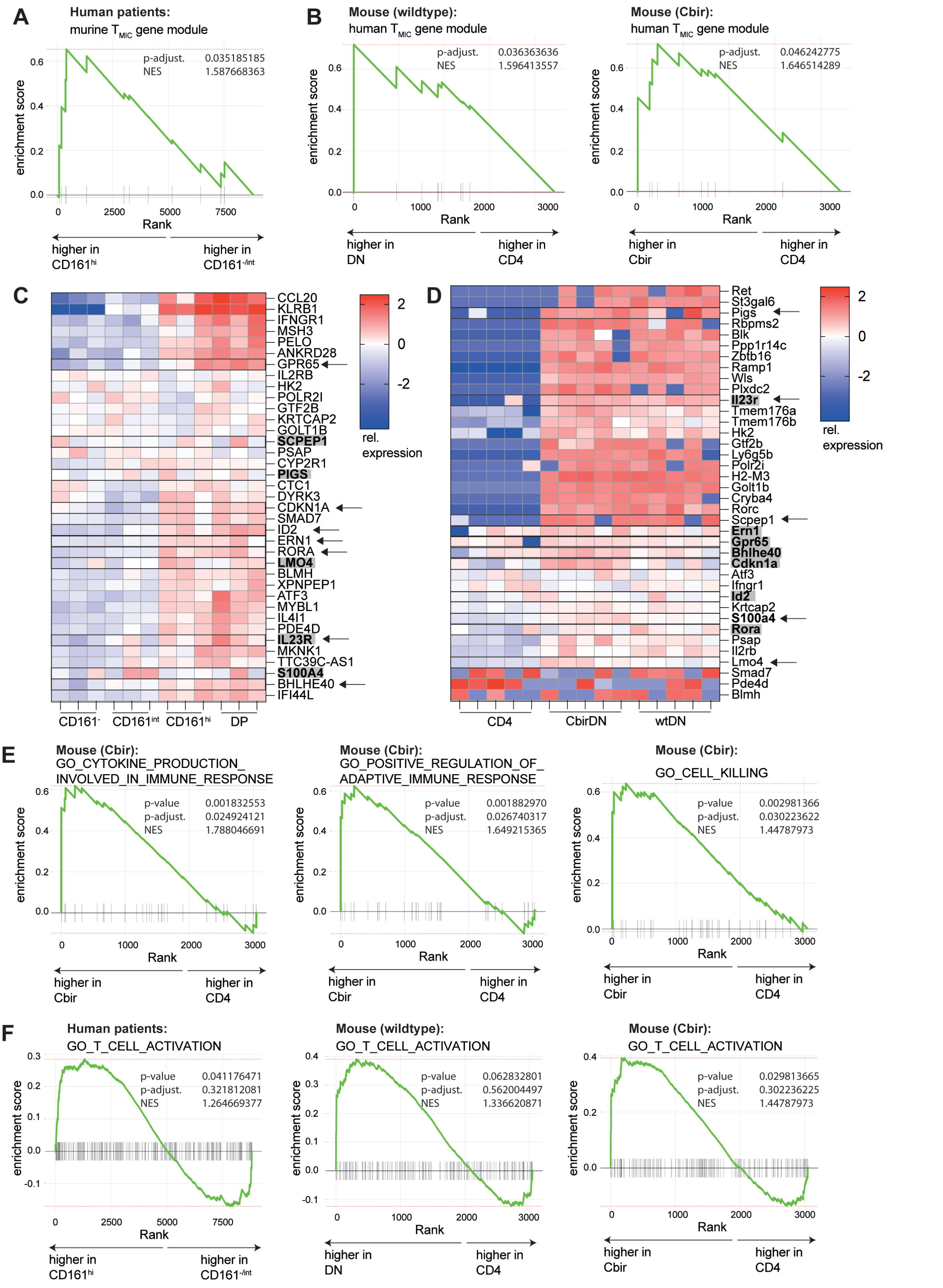


**Supplemental Figure 6: Enrichment of human and murine genes signature in the respective other species. Related to Figure 4.**

(A) Enrichment of the murine TMIC gene module (genes upregulated in murine DN cells compared to colonic CD4 T cells) in human CD161$^{hi}$ versus CD161$^{int/-}$ CD4 T cells.
(B, C) Enrichment of the human TMIC gene module (genes upregulated in human CD161$^{hi}$ compared to CD161$^{int/-}$ CD4 T cells.) in murine wildtype or Cbir DN versus colonic CD4 T cells.
(C, D) Relative expression of the combined human and murine lists of genes significantly upregulated in CD161$^{hi}$ CD4 or DN T cells. Leading edge genes driving the enrichment shown in (A-C) are marked with grey boxed while genes contributing to the enrichment in the respective other species are marked with arrows.
(E) GSEA plots depicting the enrichment of three key GO-terms in Cbir-TCR transgenic $T_{MIC}$ cells compared to murine colonic wildtype CD4 T cells.
(F) GSEA plots showing the enrichment of the top GO-term “T cell activation” in different human or murine $T_{MIC}$ cells compared to their respective control populations.

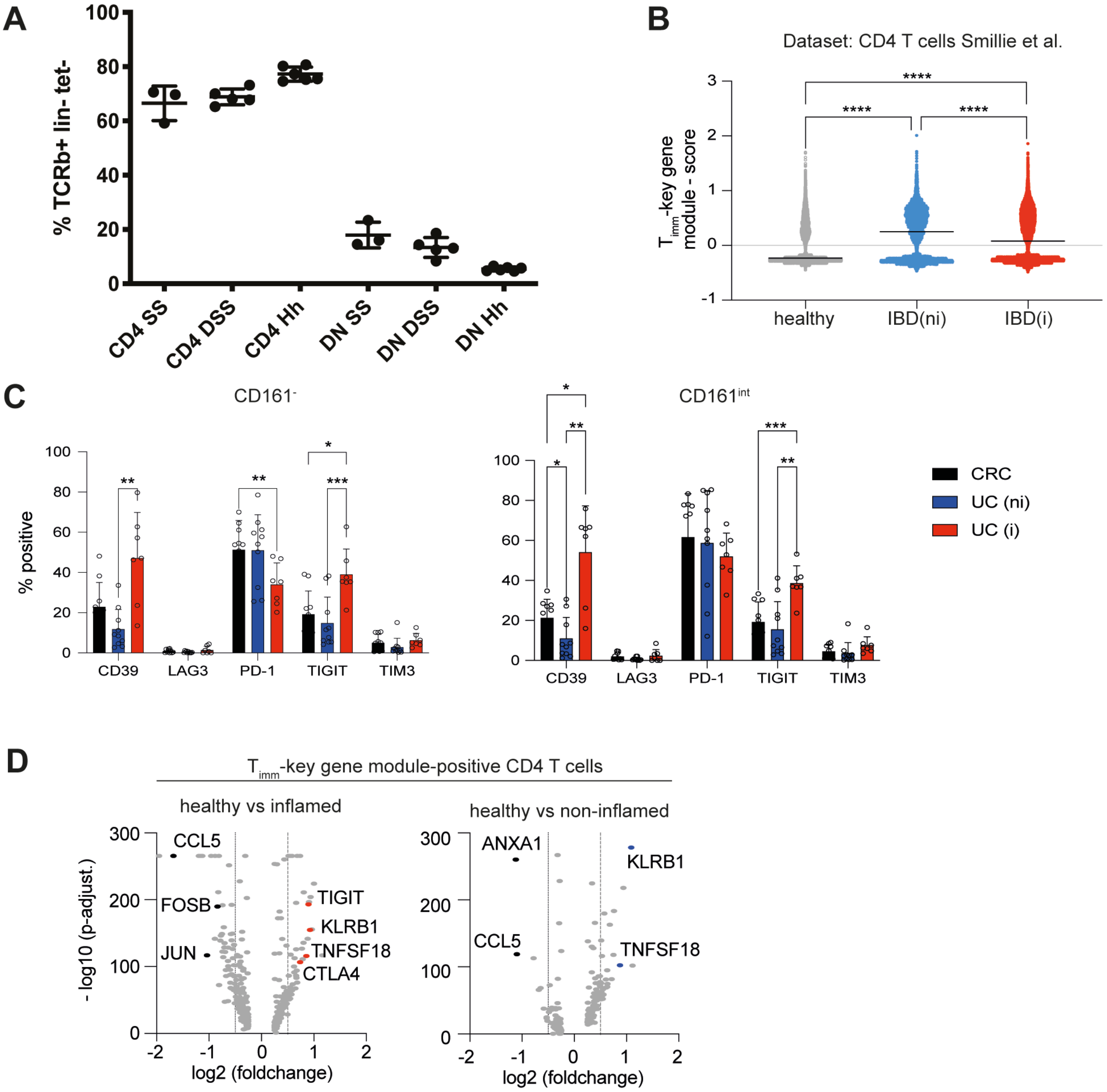


**Supplemental Figure 7: Frequencies and characteristics of human and murine $T_{MIC}$ cells in healthy and inflamed tissue. Related to Figure 5.**

(A) Quantification of CD4 and DN cell populations in steady-state, DSS, and Hh anti-IL10R driven colitis, demonstrating decrease of DN as a percent of T cells but not absence of the population at peak inflammation.

(B) Plots showing the expression score of a module of genes associated with the $T_{MIC}$ phenotype (KLRB1, ZBTB16, IL23R and IL18R1) in CD4 T cells from the study published by Smillie *et al*. The score was calculated by subtracting the average expression level of a set of randomly selected control genes from the the average expression of the module's genes on a single cell level using the *AddModuleScore* function from the R Seurat package. Module scores were compared between cells isolated from healthy controls or IBD patients with or without inflammation.

(C) Barplots summarising the expression of CD39, LAG3, PD-1, TIGIT and TIM3 on the CD161$^{-}$ or CD161$^{int}$ CD4 T cells isolated from the lamina propria of patients belonging to the indicated groups.

(D). Volcano plots showing genes differentially expressed between a subset of CD4 T cells from the indicated groups from the Smillie *et al.* dataset. Only CD4 T cells with a $T_{imm}$ -key gene module score above 0 (see B) were included in the analysis.

*p < 0.05, **p < 0.01, ***p < 0.001, ****p<0.0001; Kruskal-Wallis test with Dunn's multiple comparisons test (B), mixed effects analysis with Tukey's multiple comparisons test (C). Mean ± SEM is shown.

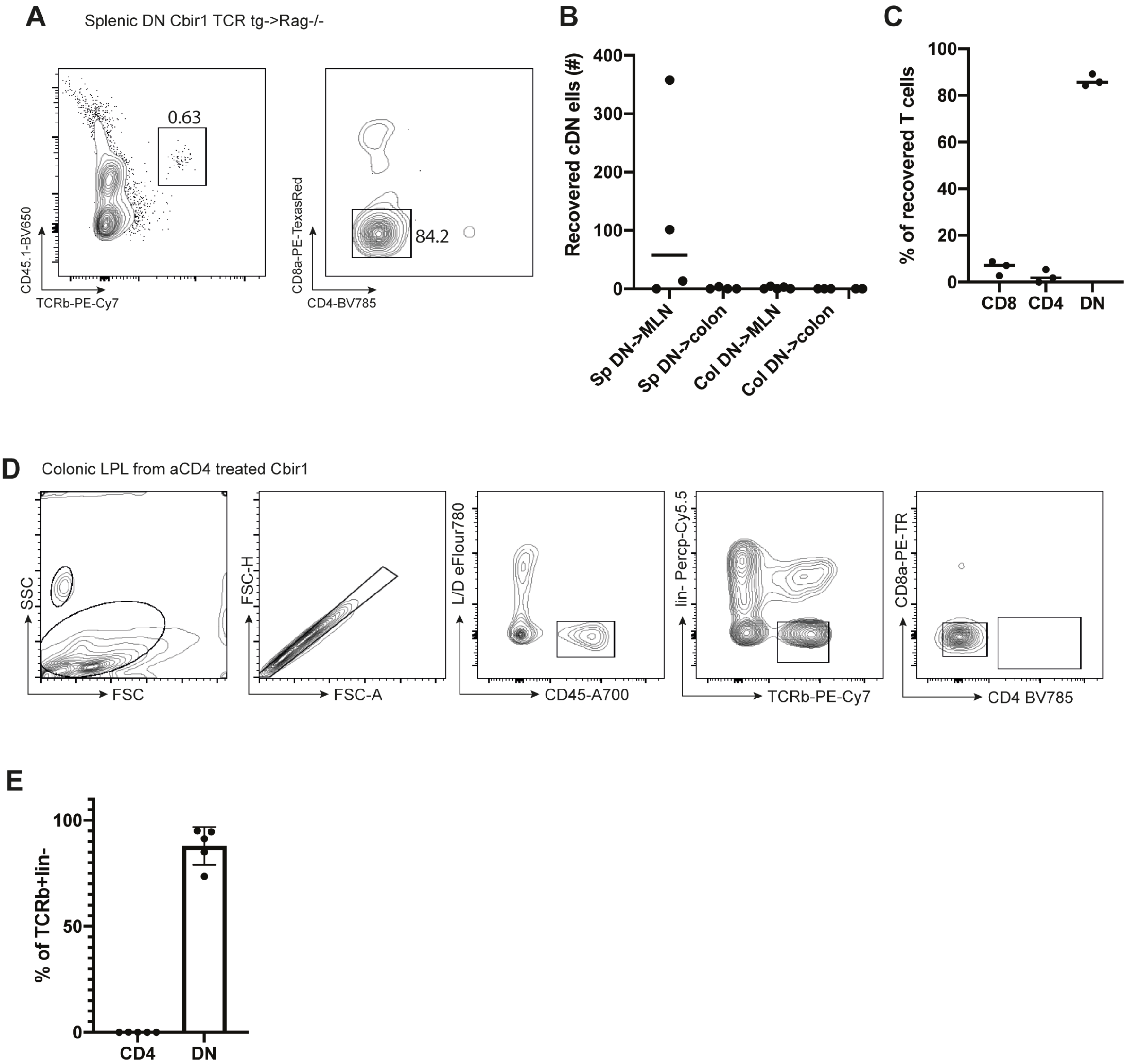


Supplemental Figure 7. Transfer of $T_{MIC}$ cells maintain DN phenotype, and aCD4 treatment depletes the CD4 subset in Cbir1 mice.

**Supplemental Figure 8: Transfer of $T_{MIC}$ cells maintain DN phenotype but do not result in colonic residency and Cbir**

(A) Representative FACS plot of MLN from a Rag-/- recipient of splenic Cbir1 DN T cells bearing the CD45.1 congenic marker (left) showing DN phenotype (right). (B) Quantification of recovered DN T cells from transfer into Rag-/-. (C) Quantification of the CD4/CD8a flow cytometric analysis of CD45.1+ T cells recovered from transfer experiment into Rag-/-. (D) Gating strategy and representative FACS of colonic T cells in aCD4 treated Cbir mice 18 days after treatment with DSS.

(E) Quantification of DN and CD4 colonic T cells in aCD4 treated Cbir mice 18 days after treatment with DSS. Data is representative of two independent experiments, n=5 per experiment.
